# Nephrectomy and high-salt diet inducing pulmonary hypertension and kidney damage by increasing Ang II concentration in rats

**DOI:** 10.1101/2024.01.25.577304

**Authors:** Qian Jiang, Qifeng Yang, Chenting Zhang, Chi Hou, Wei Hong, Min Du, Xiaoqian Shan, Xuanyi Li, Dansha Zhou, Dongmei Wen, Yuanhui Xiong, Kai Yang, Ziying Lin, Jingjing Song, Zhanjie Mo, Huazhuo Feng, Yue Xing, Xin Fu, Chunli Liu, Fang Peng, Bing Li, Wenju Lu, Jason X.-J. Yuan, Jian Wang, Yuqin Chen

**Affiliations:** State Key Laboratory of Respiratory Diseases, National Center for Respiratory Medicine, Guangdong Key Laboratory of Vascular Diseases, National Clinical Research Center for Respiratory Diseases, Guangzhou Institute of Respiratory Health, the First Affiliated Hospital of Guangzhou Medical University, GMU-GIBH Joint School of Life Sciences, Guangzhou Medical University, Guangzhou, Guangdong, China, 510120; Department of Neurology, Guangzhou Women and Children’s Medical Center, Guangzhou Medical University, Guangzhou, Guangdong, China, 510623; Department of stomatology, The First Affiliated Hospital of Guangzhou Medical University, Guangzhou Medical University, Guangzhou, Guangdong, China, 510120; Department of Critical Medicine, The Third Affiliated Hospital of Guangzhou Medical University, Guangzhou, Guangdong, China, 510150; Section of Physiology, Division of Pulmonary, Critical Care and Sleep Medicine, Department of Medicine, University of California San Diego, La Jolla, CA, USA, 92093; Guangzhou Laboratory, Guangzhou International Bio Island, Guangzhou, Guangdong, China, 510320

**Keywords:** Chronic kidney disease, Pulmonary hypertension, Renin-angiotensin-aldosterone system, Angiotensin converting enzyme 2, Metabolomics

## Abstract

**Background:** Pulmonary hypertension (PH) is a common complication in patients with chronic kidney disease (CKD), affecting prognosis. However, the pathogenesis is not clear, and the lack of a stable animal model is a significant factor.

**Methods:** In this study, a rat model of chronic kidney disease with pulmonary hypertension (CKD-PH) was developed through 5/6 nephrectomy combined with a high-salt diet. The model’s hemodynamics and pathological changes in multiple organs were dynamically assessed. Lung tissues and serum were collected from the model rats to measure the expression of ACE2, the expression levels of vascular active components related to the renin-angiotensin-aldosterone system (RAAS), and changes in the serum metabolic profile of the model.

**Results:** After 14 weeks post-surgery, the CKD-PH rat model exhibited significant changes in hemodynamic parameters indicative of pulmonary arterial hypertension, along with alterations such as right ventricular hypertrophy. However, no evidence of pulmonary vascular remodeling was observed. An imbalance in the renin-angiotensin-aldosterone system was identified in the CKD-PH rat models. Downregulation of ACE2 expression was observed in pulmonary tissues. The serum metabolic profile of the CKD-PH rat models showed distinct differences compared to the sham surgery group.

**Conclusions:** The development of pulmonary arterial hypertension in CKD-PH rats may be primarily attributed to the disruption of the renin-angiotensin-aldosterone system (RAAS), coupled with a decrease in ACE2 expression in pulmonary vascular endothelial tissues and metabolic disturbances.

## Introduction

Chronic kidney disease (CKD), a widespread global disease, has gained significant attention as a crucial public health problem in recent years, and the prevalence of this disease reported to be 10-13%^1^. Pulmonary hypertension (PH) is a common complication and a poor prognostic indicator of CKD^2, 3^. The incidence of PH in patients with CKD stage 1 and stage 5 is approximately 6% and 26.6%, respectively^4^. PH is commonly observed in 22% of patients with CKD, and it is associated with a significant risk of hospitalization and death^5^. The prevalence of PH in CKD patients without dialysis is as high as 32%^6^. The etiology and pathogenesis of CKD-related PH (CKD-PH) are largely unknown, and further studies are needed to explore the underlying mechanism of CKD-PH and develop potential treatment strategies.

Studies have confirmed that the overload of the renin-angiotensin-aldosterone system (RAAS) is involved in the pathological process of CKD, and the inhibition of RAAS activities achieves meaningful clinical benefit in patients with CKD ^7–9^. RAAS and sympathetic nervous system (SNS) play a critical role in the development of PH by modulating endothelial dysfunction and vascular remodeling^10–12^. Of note, elevation of RAAS activities was observed in PH patient, while inhibition of RAAS can ameliorate survival of PH cohorts^13, 14^. Additionally, Angiotensin II-induced migration of pulmonary arterial smooth muscle cells (PASMCs) is observed in chronic thromboembolic pulmonary hypertension (CTEPH) ^15^. A study indicated that reduced SNS activities induced by percutaneous pulmonary artery denervation (PADN) leads to downregulation of pulmonary artery pressure and promoting exercise endurance capacity ^16^. Therefore, we hypothesized that the change of RAAS and SNS activities may possibly involve in the pathophysiological process of CKD and PH. However, the mechanism behind this involvement is not fully understood.

To determine the time-course alterations of CKD and PH parameters and explore the underlying molecular mechanism of RAAS system in CKD-PH, a CKD-PH rat model will be established in this study. This study aims to investigate the role of RAAS system in the progression of CKD-PH and provide novel insights into CKD-PH molecular mechanisms regarding to develop new direction for disease management.

## METHODS

### Establishment of CKD-PH rat model

All animal experiments described in this study were conducted in compliance with the guidelines of The First Affiliated Hospital of Guangzhou Medical University Animal Care and Use Committee (approval number: 2021247). The CKD-PH animal models were established using previously established methods^17, 18^, in which a combination of 5/6 nephrectomy and a high-salt diet were applied. Specific pathogen-free male Sprague Dawley rats weighing 280±20g(8-9 weeks old) were obtained from the Guangdong Provincial Medical Experimental Animal Centre and housed in the SPF grade animal room of the State Key Laboratory of Respiratory Disease, Guangzhou Medical University. As illustrated in Figure 1, the rats were randomly divided into sham and 5/6 nephrectomy groups at 8 weeks and 14 weeks. The rats were anesthetized with isoflurane (3% for induction, then 2.5% for maintenance). Nephrectomy was performed after shaving and disinfecting the surgical area with 75% alcohol. A 1.5-cm incision was made through the skin, subcutaneous tissue, and muscles to expose the left kidney, and the upper and lower poles of the left kidney were removed. One week after the surgery, the right kidney was completely removed. In the sham group, the same incision was made without nephrectomy. Seven days after the surgery, rats in the 5/6 nephrectomy group were fed a high-salt diet (8%), while rats in the sham group received a normal-salt diet (0.4%).

**Figure 1.**
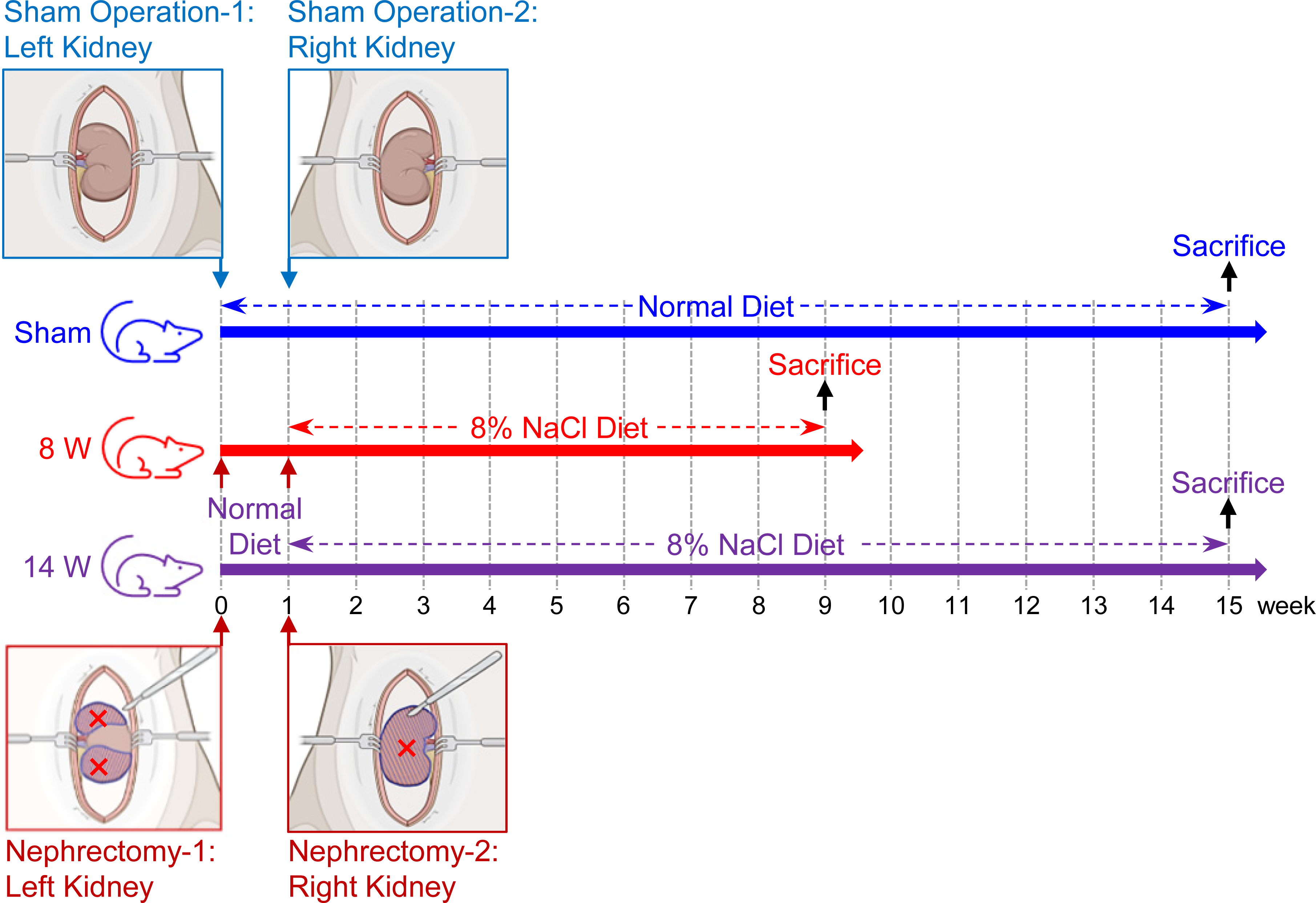
Protocol for the establishment of CKD-PH rat model.

### Hemodynamic and histological measurements

After anesthesia, hemodynamic parameters were measured in rats from each group using a BIOPAC MP150 data acquisition system (BIOPAC systems, Inc., Santa Barbara, CA), including right ventricular systolic pressure (RVSP), systolic blood pressure (SBP), and diastolic blood pressure (DBP). The ratio of RV mass to the mass of left ventricle plus septum (RV/(LV+S)) was calculated to assess right ventricular hypertrophy (RVH). In addition, the kidney, liver, and spleen were weighed. The lung, kidney, and heart were then formalin-fixed, paraffin-embedded, and stained with H&E and Masson’s trichrome (KGMST-8003) to examine any pathological changes in these tissues. Pulmonary vascular remodeling was evaluated by determining the ratio of wall thickness to total vascular diameter (WT%) and the number of vessels per square millimeter (mm^2^).

### Immunofluorescent staining

To determine the expression of ACE2, CD31, and α-SMA, immunofluorescent staining of lung tissue was performed using previously established methods^19^. After dewaxing, dehydrating, and antigen retrieval, lung tissue slides were incubated with primary antibodies, followed by secondary fluorescent antibodies. The expression of ACE2, CD31, and α-SMA was quantified using case viewer. The primary antibodies used were ACE2 (Abmart, Cat.#T55787M), CD31 (MLBIO, Cat.#ml058803), and α-SMA (Sigma-Aldrich, Cat. #A5228).

### Echocardiographic assessment

Stroke volume, cardiac output (CO), ejection fraction(EF) and pulmonary acceleration time/pulmonary ejection time (PAT/PET) were measured with the same protocols as described previously^20^.

### Isometric tension measurement of isolated pulmonary artery

Isolated pulmonary artery (PA) rings were prepared according to previously described methodology^21, 22^. The PA rings were allowed equilibrium and optimally stretching in Krebs-Henseleit solution (containing NaCl 118.0 mM, NaHCO_3_ 25.0 mM, KCl 4.7 mM, NaH_2_PO_4_ 1.2 mM, CaCl_2_ 1.8 mM, MgSO_4_ 1.2 mM and glucose 11.1 mM) that was bubbled with 95% O_2_ and 5% CO_2_ for 100 minutes. To confirm the viability of the PA rings, KCl (60 mM) was used to induced contraction. Once the PA rings reached maximum amplitude, Ach (10μM) was applied to determine the integrity of the endothelium. Phenylephrine (1 μM) was then employed to achieve optimal contraction in calcium-free solution. Then, PA rings were allowed to restore to their basal tension after Phenylephrine-induced contraction. To evaluate the concentration of AngⅡin plasma, PA rings were incubated with plasma from both sham group and 5/6 nephrectomy groups, using AngⅡ as a positive control.

### Metabolomics analysis

Blood was collected from the caudal vein using 5ml EDTA tubes. The plasma was obtained through low-temperature centrifugation at 5000 g for 10 minutes. To investigate the plasma metabolomics profile of the CKD-PH rat model, LC–MS/MS technique was used in a high-resolution mass spectrometer Q Exactive HF system (Thermo Fisher Scientific, USA). Data was collected and further analyzed using the R 4.1.1 DeSeq2 package. Significant differential metabolites were identified through univariate analysis, based on ∣log2 fold change (FC)∣>1 and P < 0.05 (sham vs 5/6 Nx group). The edgeR and limma algorithms were also used to select more reliable differential metabolites. The differential metabolites were integrated and ranked using robust rank aggregation (RRA). The candidate metabolites were analyzed and annotated using Kyoto Encyclopedia of Genes and Genomes (KEGG) databases (http://www.genome.jp/kegg/). The raw metabolomics data for this study is available in the Supplemental Materials.

### Measurements of plasma renin, Ang II, aldosterone and Ang (1-7)

Plasma samples were prepared as described above. The listed ELISA kits (Renin, MEIMIAN Co.Ltd, Beijing, China, Cat.#: MM-034R2; AngII, MLBIO, Cat.#: ml058803; Aldosterone, MEIMIAN Co.Ltd, Beijing, China, Cat.#: MM-0555R2; and Ang(1-7), Shanghai Jianglai industrial Limited By Share Ltd, China, Cat.#: JL21363-96T) were used to measure the concentrations of renin, Ang II, aldosterone, and Ang(1-7) following the manufacturer’s instructions.

### Western Blot

To obtain whole-tissue extracts, lysed in RIPA lysis buffer (50 mmol/L Tris pH 7.4, 150 mmol/L NaCl, 1% NP-40, 0.5% sodium deoxycholate, and 0.1% SDS) supplemented with PMSF ( 100mmol/L GBCBIO), protease inhibitor cocktail (E2, Roche) and phosphatase inhibitor cocktail (E3, Sigma) (RIPA:PMSF:E2:E3=100:2:1:1). After three repeated grind, tissue extracts were cleared by centrifugation at 4°C. Tissue extracts were quantified using Bradford assay, and equal amounts of soluble proteins were mixed with SDS sample buffer, boiled, separated by SDS-PAGE, and blotted onto polyvinylidene difluoride membranes (Millipore). Membranes were then blocked for 1 hour with 5% dry milk in PBST (PBS supplemented with 0.1% Tween-20) buffer and incubated overnight at 4°C with primary antibody (anti-ACE2, Abcam; anti-Actin, Abcam) diluted in PBST containing 1% BSA (Sigma). After three washes with PBST, membranes were incubated with HRP-conjugated anti-mouse or anti-rabbit IgG. Membranes were washed five times with PBST, and the blotted protein bands were revealed by the ECL system (Millipore).

### Statistical analysis

In this study, “n” refers to the number of animals, and data is presented as means ± standard deviations (SD). We used one-way ANOVA with Tukey’s post hoc analysis for multiple group comparisons and student’s t-test for two-group comparisons. In cases where the data deviated significantly from normality, we applied non-parametric tests (Mann-Whitney test). Receiver-operating-characteristic (ROC) curve analysis was used to assess the diagnostic accuracy of candidate metabolites. Statistical analysis of metabolomics data was performed using the R 4.1.1 DeSeq2 package. A statistical significance level of \**P*<0.05 and \*\**P*<0.01 was considered significant.

## Results

### 5/6 Nx and high-salt diet induced progression of chronic kidney disease

In comparison to the sham group, the levels of serum creatinine and urea nitrogen gradually increased in the 5/6 Nx group with the passage of time, indicating impaired renal function (Figure 2A). However, the total urine protein level of 24 hours in the 5/6 Nx 8-week group was much higher than that in the sham group, and slightly higher than that in the 5/6 Nx 14-week group (Figure 2A). Additionally, Figure 2C revealed remarkable swelling and sclerosing glomeruli, dilated renal tubules, and marked interstitial fibrosis in the 5/6 Nx-induced 8-week group. As a result, prolonged 5/6 Nx and high-salt diet treatment forced the swelling and sclerosing glomeruli slowly converted to normal size with structural derangement. Simultaneously, renal tubules gradually developed with an enlarged inner lumen accompanied by obvious renal interstitial fibrosis.

**Figure 2.**
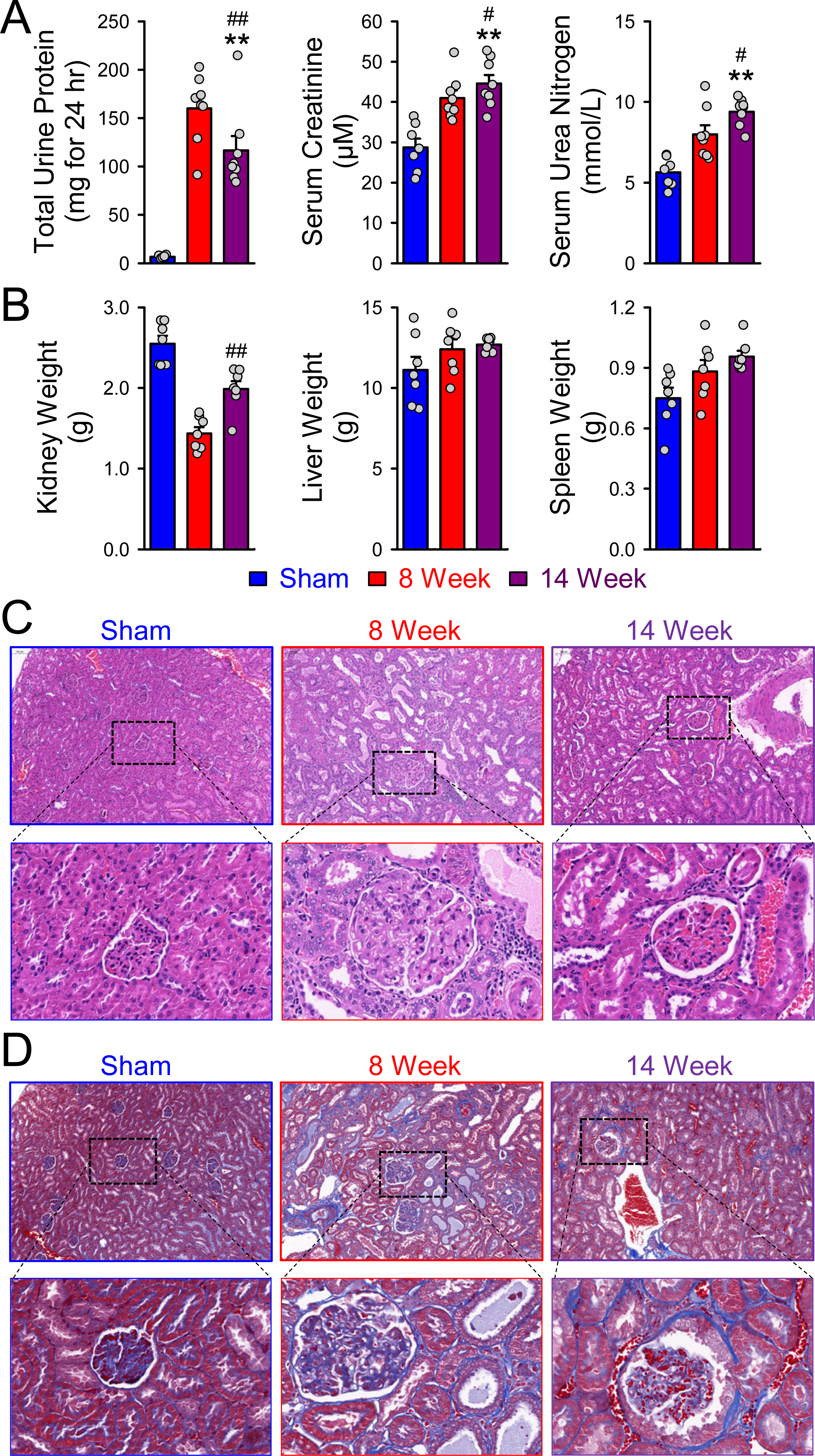
Progressive renal dysfunction and accelerated collagen deposition were confirmed in 5/6 Nx and high-salt diet-treated rats. A Bar graphs presenting the total urine protein, serum creatinine and serum urea nitrogen. B bar graphs showing weight changes of kidney, liver and spleen during modeling. (C-D) Representative images of HE(C) and Masson’s(D) trichrome staining revealing renal structural modification and collagen deposition in the kidney sections from each group of rats. Data shown are means ± SEM; n = 7 in each group. * *P < 0.05*, * \**P < 0.01,* significantly different as indicated.

To determine the extent of renal fibrosis, Masson’s trichrome staining was conducted, in which accumulation of collagen presented as blue staining and fibrin showed as red staining. As shown in Figure 2D, the region of blue staining gradually expanded following the 5/6 Nx and high-salt diet treatment for 8 and 14 weeks, indicating the progression of renal fibrosis. In the process of the 14-week modeling, the weight of the liver and spleen showed no significance between groups (Figure 2B). However, the weight of the kidney in the 5/6 Nx 8-week group was significantly lower than that in the sham group, and remarkably recovered in the 5/6 Nx 14-week group (Figure 2B) (*P*<0.05).

### 5/6 Nx in combination with high-salt diet provoked pulmonary hypertension and impairment of heart function

In this study, typical hemodynamic indexes of PH were evaluated. As shown in Figure 3A-B, We found that a gradual increase over time in both RVSP and the RV/(LV + S) ratio, indicating the establishment of CKD-induced PH rat models. Furthermore, 5/6 Nx-14 weeks group exhibited significantly elevated RVSP and RV/ (LV + S) ratios in comparison to the sham group (*P*<0.05). Raised systemic blood pressure ( mean arterial blood pressure(mABP), SBP, and DBP) with significant difference was observed in the 5/6 Nx groups compared to the sham group (Figure 3C, *P*<0.05).

**Figure 3.**
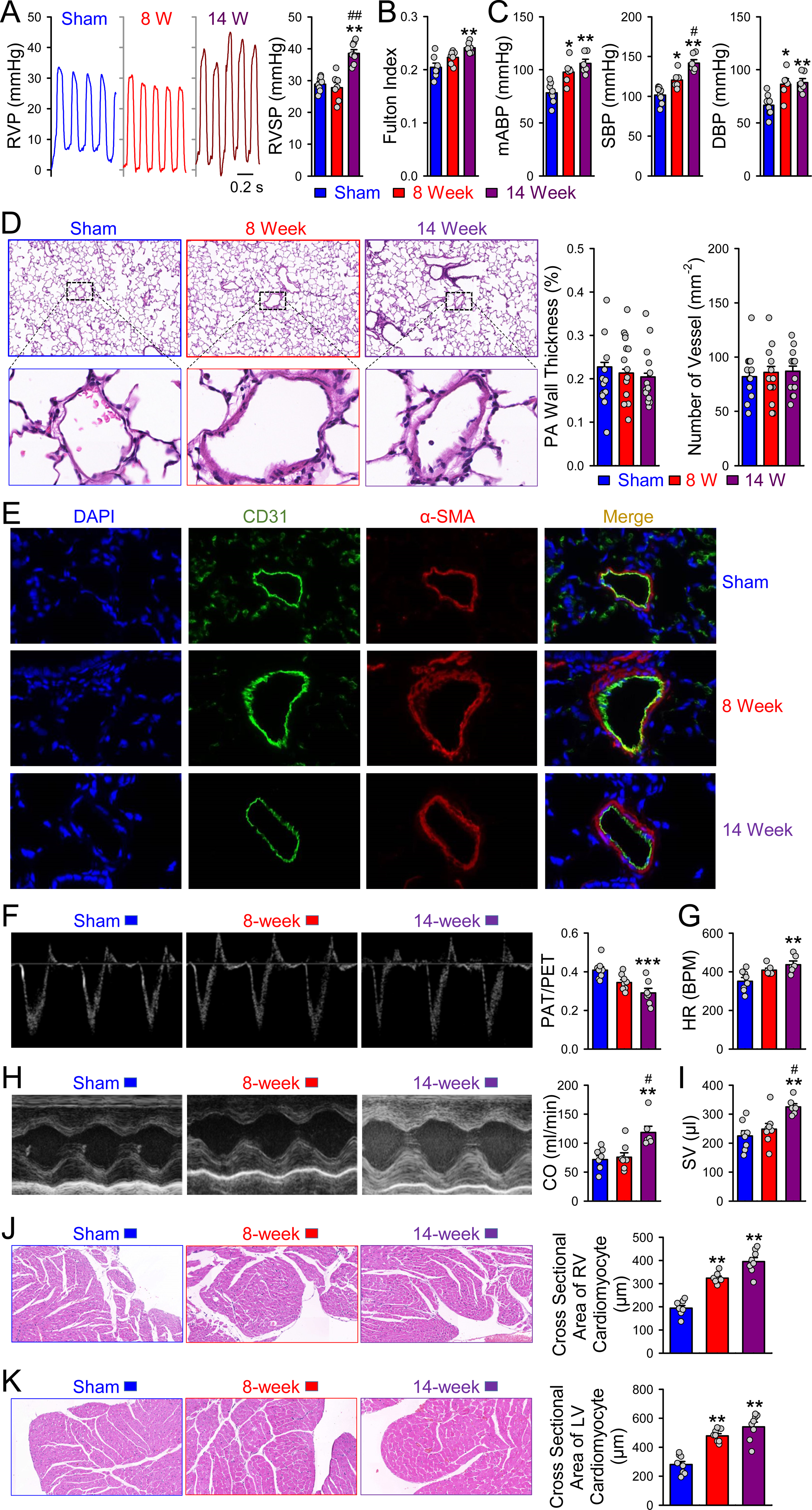
Increased hemodynamic measurements and right ventricular hypertrophy without pulmonary vascular remodeling was identified in 5/6 Nx and high-salt diet-treated rats. Representative traces and bar graph (A) of the right ventricular systolic pressure (RVSP). Bar graph (B) finding the Fulton index [RV/(LV + S)] of sham and 5/6 Nx and high-salt diet-treated rats. Bar graphs of systemic blood pressures, including mABP, SBP and DBP were presenting in C. (D) Staining with HE shows the histological and pathological changes in the pulmonary arteries in the lung sections of each group; analyzed data presents wall thickness (WT (%)) and vessel numbers per mm^2^ of the pulmonary arteries in each group of rats. Typical immunofluorescence staining images (E) in lung sections from rats of each group. Staining of blue, green and red is represented as Dapi, CD31 and α-SMA, respectively. Scale bar represents 20 μm as indicated. Representative images and summary data of pulmonary acceleration time/pulmonary ejection time (PAT/PET) and stroke volume (SV) are shown in F and I, respectively. Bar graphs of cardiac output (CO) and heartbeat (HR) from each group are presented in G and H. H&E staining of heart slides showing right and left myocardial cells show in J and K. Summarized data showing cross sectional area of right and left myocardial cells among the sham, 5/6 Nx-8-week group and 5/6 Nx-14-week group. Data shown are means ± SEM; n = 7 in each group. * *P < 0.05*, * \**P < 0.01,* significantly different as indicated.

Interestingly, unlike other PH animal models, we did not observe significant smooth muscle media wall thickness and distal pulmonary vascular remodeling in the 5/6 Nx and high-salt diet-induced CKD-PH rat models. This was demonstrated by the percentage of wall thickness (WT%) and vessel numbers per mm2, respectively (Figure 3D). Immunofluorescent double staining of α-SMA (red) and CD31 (green) showed no significant difference in the thickness of pulmonary artery smooth muscle medial wall between groups (Figure 3E).

Cardiac function was assessed through hemodynamic indexes, echocardiographic analysis, and pathological examination. Left heart function parameters, including left ventricular EF, cardiac output (CO), and stroke volume (SV), as well as right heart function measurements, such as PAT/PET, were detected by echocardiography. SV, HR, and CO were remarkably higher in the 5/6 Nx-14-week group than in the sham group and 5/6 Nx-8-week group (*P*<0.05), as shown in Figure 3G-I. In addition, as illustrated in Figure 3F, it is important to highlight that PAT/PET exhibited a notably lower value in the 5/6 Nx-14 weeks group than in the sham group (*P*<0.05).

Right and left heart myocardial hypertrophy in rats treated with the combination of 5/6 Nx and high-salt diet throughout the 8- to 14-week time course was evaluated by cardiac histology, which was manifested by a significant increase in the cross-sectional area of myocardial cells (*P*<0.05), as depicted in Figure 3J-K.

### Disorder of renin-angiolensin-aldosterone system was found in CKD-PH rat models

We observed a strong upward trend in the plasma concentration of AngⅡ, while the levels of Ang (1-7) decreased steadily during the 5/6 Nx+ high-salt diet treatment (*P*<0.05) (Figure 4A). Renin and aldosterone (ALD) levels were significantly elevated in the 5/6 Nx+ high-salt diet 8-week and 14-week groups, compared to the sham group. However, the levels of renin and aldosterone (ALD) were slightly lower in the 8-week group than in the 14-week group (Figure 4A). To further determine the effect of the RAAS system on pulmonary vascular tone, we evaluated the isometric tension of isolated PAs. We observed that plasma from the 14-week 5/6 Nx groups significantly stimulated pulmonary vasoconstriction in PA rings, whereas such phenomenon was not seen in the plasma of the sham group (Figure 4C, E). However, losartan (a suppressor of AngⅡ) dramatically blocked the 14-week plasma-induced constriction of PA rings (*P*<0.05). As indicated in Figure 4F, G, ACE2 was highly expressed in endothelial cells, whereas significant reduction of ACE2 was observed in the treatment of 5/6 Nx and high-salt diet, suggesting that ACE2 was suppressed in the CKD-PH rat model. Similarly, Western blot analyses on ACE2 in the lung border tissue (mainly containing lung EC) also confirmed the downregulation of ACE2 in the 5/6 Nx-8-week and Nx-14-week group than the sham group (Figure 4H).

**Figure 4.**
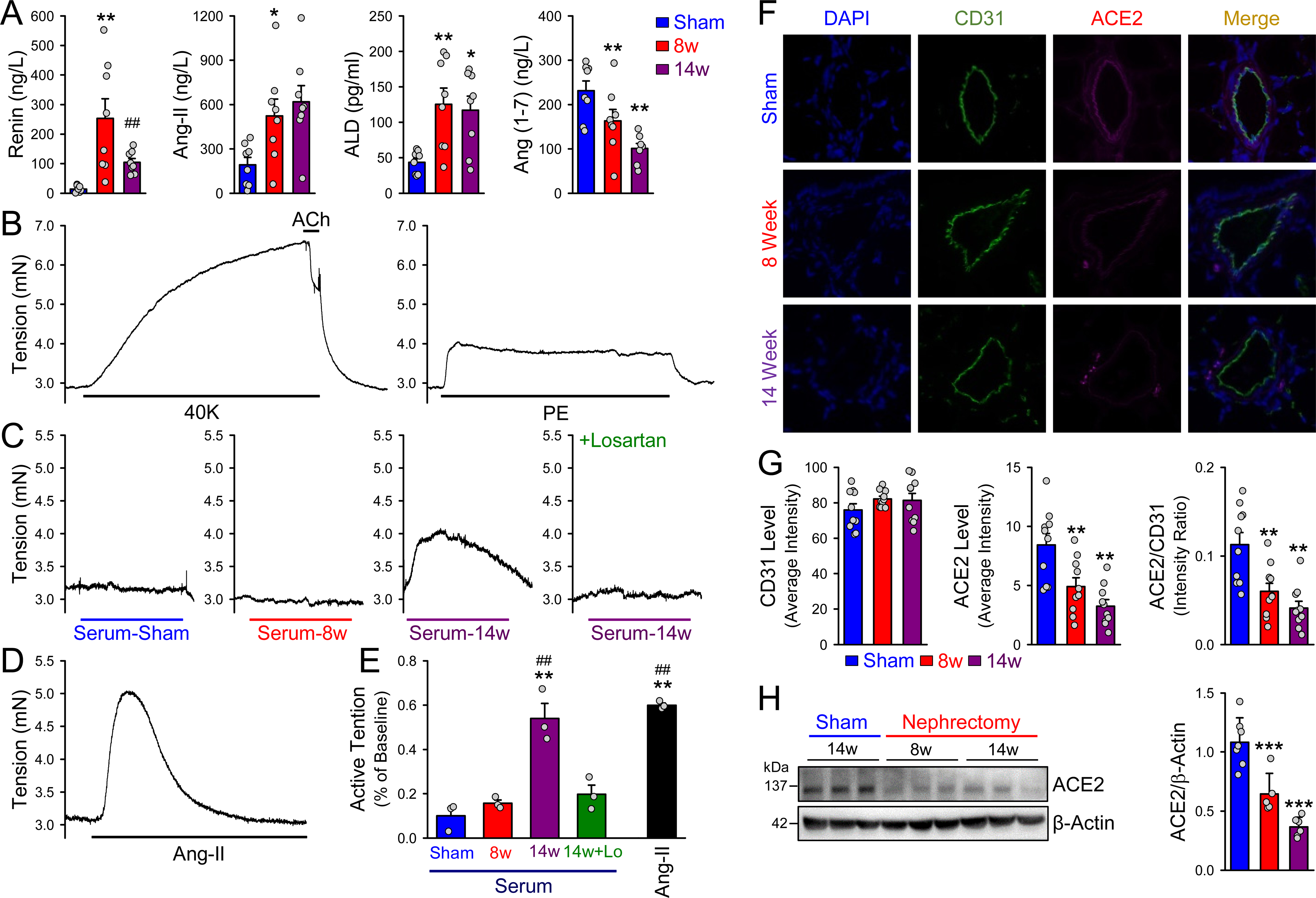
Impairment of RASS in CKD-PH rat model. Bar graphs of renin, AngⅡ, aldosterone (ALD) and Ang (1-7) concentration from each group present in A (mean ± SEM; n = 8 in each group). Representative curves (B-D) presenting plasma/AngⅡ-mediate contraction in isolated pulmonary artery (PA) rings precontracted with PE. (E)Summarized data (mean ± SEM; n = 3 in each group) showing the contraction of isolated PA rings in the presence of plasma or AngⅡ treatment. Typical immuno-fluorescence staining images (F) and summary data (mean ± SEM; n = 6 in each group) of average immunofluorescent intensity for CD31, ACE2 and ACE2/CD31 (G) in lung sections from rats of each group. Staining of blue, green and red is represented as Dapi, CD31 and ACE2, respectively. Scale bar represents 20 μm as indicated. Expression of ACE2 in lung tissue were measured by western blot of the sham, 5/6Nx-8-week group and 5/6Nx-14-week group. n=5-7. * *P* < 0.05, * \**P* < 0.01, significantly different as indicated.

### Identification of potential differential metabolites

Untargeted metabolomics was used to detect and analyze the plasma metabolites of the different groups. A total of 12,819 metabolites were detected and 2,092 differential metabolites were identified based on a |log2 FC|>1 and P<0.05 cutoff. Among the differential metabolites, 1,460 metabolites were up-regulated (represented as pink plot) and 632 metabolites were down-regulated (represented as blue plot) in the 5/6 Nx-operated groups compared to the sham group (Figure 5A). Using edge R and limma packages in R, 1,070 and 972 up-regulated differential metabolites were also identified (Figure 5B-C, represented as pink plot). These differential metabolites were integrated and ranked with RRA, and 400 up-regulated metabolites were detected in the operated groups (Figure 5D). A Venn diagram analysis indicated that 377 common up-regulated differential metabolites were detected while using different algorithms (Figure 5E). Based on KEGG database, 17 up-regulated differential metabolites were found (Figure 5F). The results of this analysis provide a comprehensive and detailed landscape of the metabolic changes associated with the development of CKD-induced PH.

**Figure 5.**
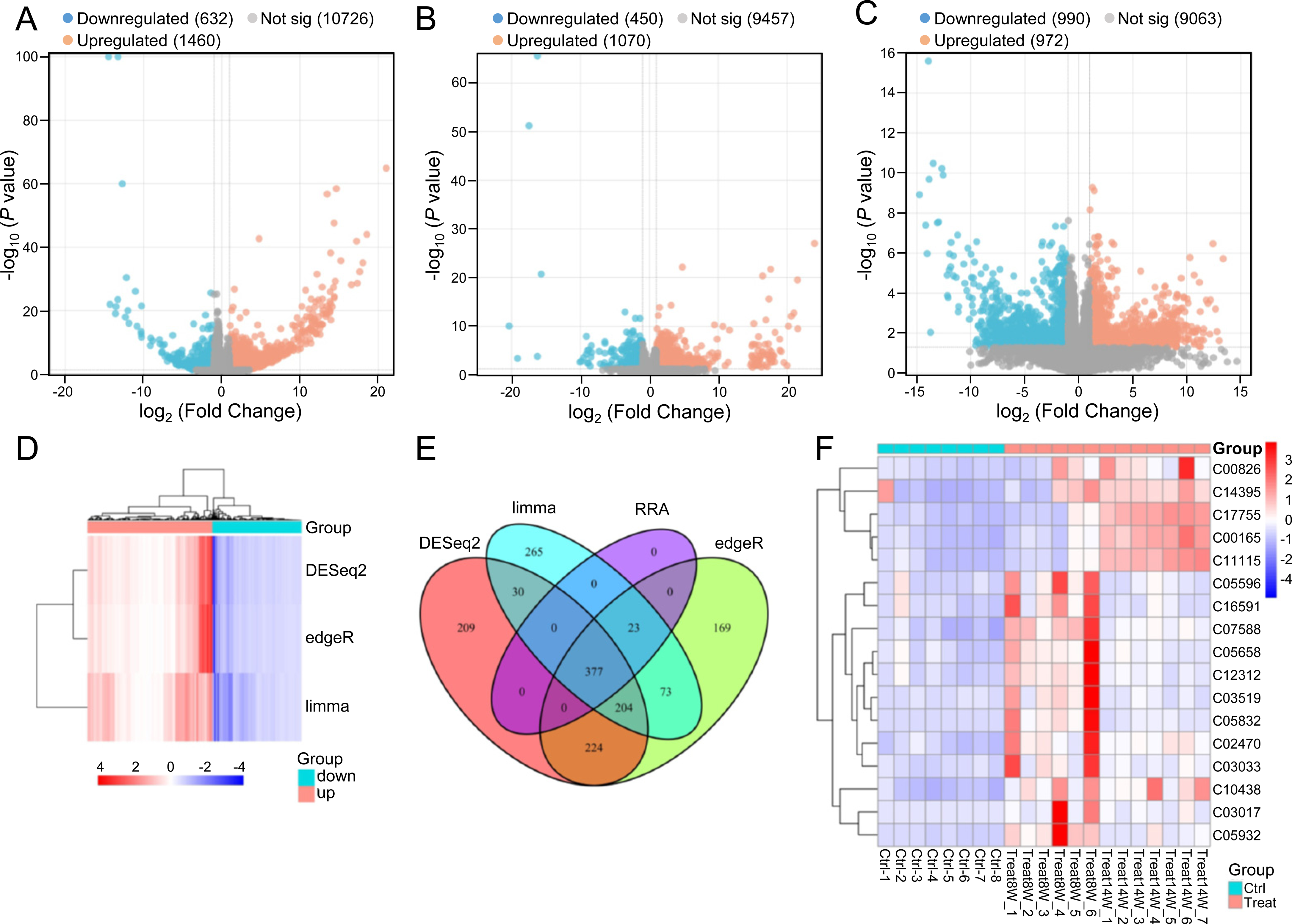
Potential differential metabolites were confirmed in rats treated with 5/6 Nx and high-salt diet. Volcano Plots data obtained from R analysis using deSeq2, edgeR and limma package showing differential metabolites among sham group and 5/6 Nx plus high-salt diet induced groups (A-C). Heatmap (D) presenting differential metabolites between sham and operated groups. Venn chart indicating common increased differential metabolites, measured by different algorithm (E). The identified 17 up-regulated differential metabolites obtained from KEEG database were recorded, showing in heatmap (F).

### Functional analysis of the identified differential metabolites

The cluster analysis allows us to identified 17 differential metabolites which were dramatically increased in the treated groups (Figure 6A). As illustrated in Figure 6B, strong correlations were observed between these metabolites. Significant negative correlations were confirmed between C00165, C11115, C17755 and other metabolites (coef<0). Advanced analysis presented the expression of these 17 metabolites in sham and treated groups (Figure 6C). Table 1 demonstrated the corresponding KEGG IDs of the 17 metabolites mentioned above. Non-parametric tests verified statistical significance between sham and treated groups for the top five up-regulated metabolites (C00165, C00836, C11115, C17755 and C05932) (*P*<0.01, Figure 6D). Additionally, all these five metabolites yielded significant AUC-ROCs (AUC>0.85, *P*<0.05) in discriminating sham from treated groups (Figure 6F). Based on KEGG database, eighty-three signaling pathways related to these five metabolites were involved, including the RAAS (aldosterone synthesis and secretion). Based on time-series analysis, all the metabolites were divided into 4 clusters (Figure 6E). Interestingly, C00165 (diacylglycerol, DAG) was included in cluster 1, indicating a consistent accumulation of C00165 continuously over the observed time period (Figure 6E). Kruskal-Wallis test was then performed to further prove this phenomenon (Figure 6G).

**Table1.**

| KEGG_ID | Name of Metabolites |
| --- | --- |
| C00165 | Diacylglycerol |
| C00836 | Sphinganine |
| C02470 | Xanthurenic acid |
| C03017 | O-Propanoylcarnitine |
| C03033 | beta-D-Glucuronoside |
| C03519 | N-Acetyl-L-phenylalanine |
| C05596 | p-Hydroxyphenylacetyl glycine |
| C05658 | Indoxyl |
| C05832 | 5-Hydroxyindoleacetyl glycine |
| C05932 | N-Succinyl-L-glutamate 5-semialdehyde |
| C07588 | Salicyluric acid |
| C10438 | Cinnamic acid |
| C11115 | 5-Methyl-2-furaldehyde |
| C12312 | Indolin-2-one |
| C14395 | alpha-Methylstyrene |
| C16591 | 3-Carbamoyl-2-phenylpropionic acid |
| C17755 | Dopamine quinone |

**Figure 6.**
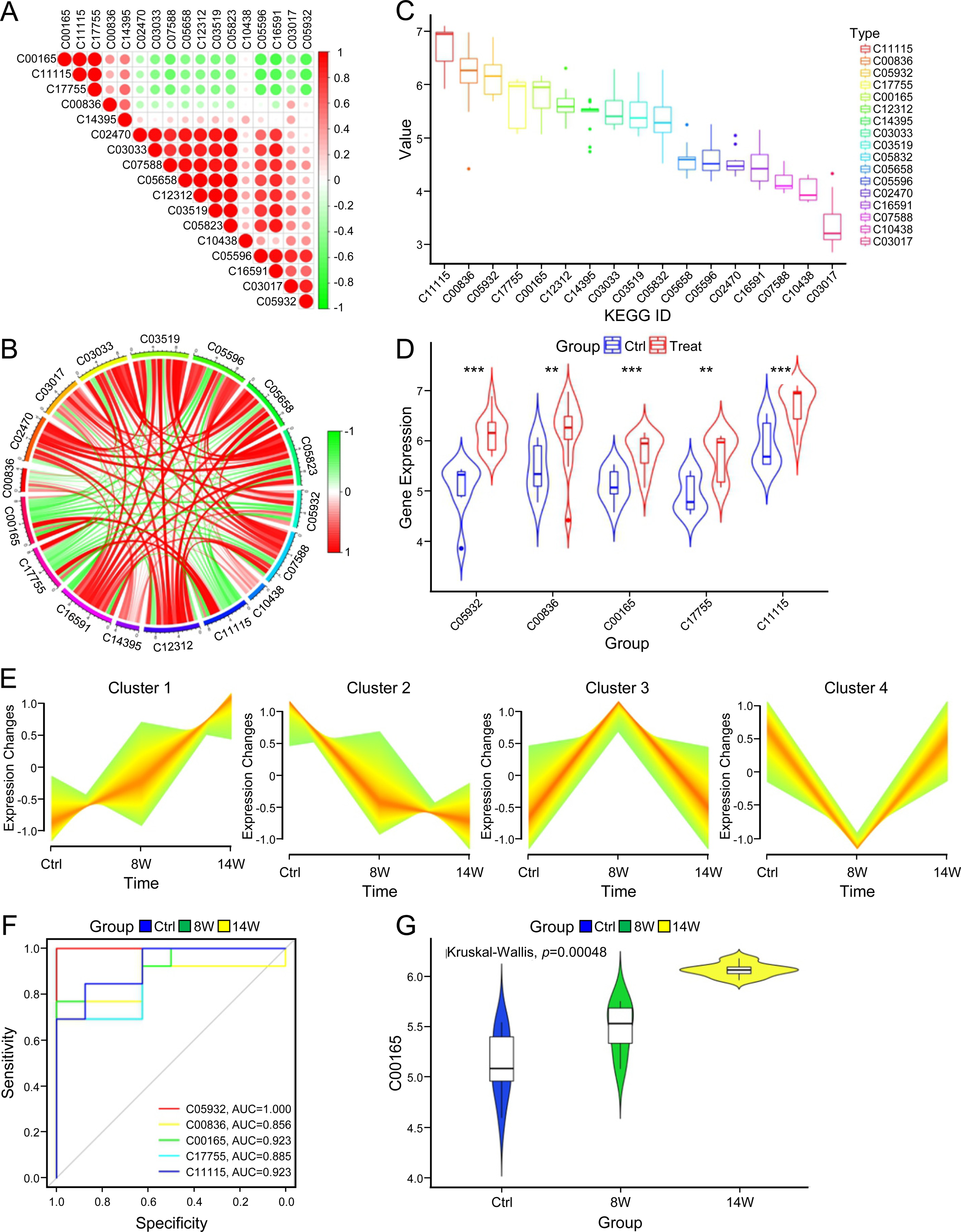
Further analysis of identified differential metabolites. Correlations of these identified differential metabolites were showing in A and B. Box plot (C) revealing relative expression of identified differential metabolites among control and treated group. Violin plot (D) suggesting relative expression of five top identified differential metabolites between sham and treated group. Time-varied metabolites were divided into four clusters in accordance with time-series trend analysis (E). ROC (F) verified diagnostic reliability of five top identified differential metabolites. The Kruskal-Wallis analysis (G) illustrates a time-dependent increase in metabolite C00165.

## Discussion

Herein, we successfully developed a rat model of CKD-PH through the combination treatment of 5/6 nephrectomy and high-salt diet. Firstly, we identified typical changes of CKD, such as an increased expression of urine protein, urea nitrogen, and creatinine as well as classic pathological alterations, which were consistent with previous studies^23–25^. Secondly, we observed characteristic hemodynamic indexes of PH (e.g., elevation in RVSP and RVH) in this animal model. However, unlike other subtypes of PH, classic pulmonary vascular remodeling was absent, indicating a different pathogenesis of CKD-PH. Thirdly, we confirmed impairment of heart function characterized by decrease in PAT/PET, an increase in blood pressure, HR, SV, and CO, and enlargement of myocardial cells in this CKD-PH animal model. Despite from the findings mentioned above, our results elicited persistent enhancement of RAAS in the CKD-PH rat model, ultimately leading to constriction of the pulmonary artery. To further explore the molecular mechanism of RAAS in CKD-PH, we performed untargeted metabolomics and identified 17 upregulated differential metabolites, among which DAG was selected based on KEEG database. Notably, we highlighted that the elevated levels of DAG induced by the RAAS may possibly played a crucial role in the development of CKD-PH.

In recent years, there has been growing interest in CKD-PH due to its high prevalence ^4^. However, little is known about the development of animal models for CKD-PH. Classic rodent models, such as 5/6 nephrectomy-induced CKD ^23^, adenine-induced CKD^26^, high-salt diet-induced CKD^27^, and 5/6 nephrectomy and high-salt diet-induced CKD^28^, have been frequently used in earlier study. Moreover, activation of the RAAS system has been shown to lead to high blood pressure and cardiac remodeling (myocardial hypertrophy and fibrosis) in CKD patients and animal models^29–33^.

In this study, we employed 5/6 nephrectomy and high-salt diet intervention to generate a rat model of CKD-PH. After conducting 14 weeks of modeling, we identified hallmark characteristics of CKD, including increased urine protein, serum creatinine and urea nitrogen levels, and distinctive pathological changes within the kidney. Additionally, we also obserevd indication of PH, namely enlarged RVSP and RVH, and cardiac remodeling. These findings suggested that we successfully established a CKD-PH rat model. Nonetheless, vascular remodeling, a prominent and characteristic pathological change of PH, is notably absent in this model^34, 35^, indicating that the pathogenesis of CKD-PH may differ from other PH subtypes and its underlying mechanism requires further investigation.

RAAS is a cascade of peptide products that regulate blood pressure, body fluid, and electrolyte balance. Evidence from both experimental animal models and human cohort studies with chronic kidney disease (CKD) has shown that an imbalance in RAAS is involved in the development and progression of CKD^8, 36–38^. Inhibition of RAAS has been proven to substantially suppress the development of CKD^39–43^. Intriguingly, studies have shown the involvement of RAAS in the development of PH^44^. Increased expression levels of renin, angiotensin I and II, and angiotensin converting enzyme (ACE) have been observed in patients with idiopathic pulmonary arterial hypertension (IPAH)^13^. Collective studies has reported that RAAS blockers can ameliorate monocrotaline-induced upregulation of RAAS and right ventricular hypertrophy^13, 45, 46^. A previous study indicated that Neprilysin inhibitor sacubitril and the angiotensin receptor blocker valsartan dramatically improved pulmonary artery pressure in patients with heart failure and reduced ejection fraction^47^. The ACE inhibitor, Captopril, has been proven to improve prognosis for patients with chronic obstructive pulmonary disease (COPD)-related PH^48, 49^. It is worth mentioning that increased expression of renin and angiotensin II levels were reported in patients withCTEPH, which eventually leads to the enhancement of migration of pulmonary arterial smooth muscle cells (PASMCs)^15^. However, of these studies to date the role of RAAS in CKD-PH remains largely unknown. By establishing a rat model of CKD-PH with 5/6 nephrectomy and a high-salt diet, we found an imbalance of RAAS (upregulated renin, aldosterone, and angiotensin II, and downregulated Ang1-7 and ACE2) in the CKD-PH rat model, indicating that the development of CKD-PH requires essential RAAS associated activities. The RAS is intricately balanced between the detrimental ACE1-Ang II-AT1R axis and the beneficial ACE2-Ang-(1– 7)-Mas axis. Suppressed ACE2 or aggra-vated ACE1 have been implicated in PH^50^. The ACE1-Ang II-AT1R arm promotes vasoconstriction, proliferation, and growth, which would be counteracted by the ACE2-Ang-(1–7)-Mas arm^51^. We found that We found a decreased expression of ACE2 in the pulmonary vascular endothelium and lung tissue in the CKD-PH rat model, which may lead to an inability to counterbalance the continuous enhancement of the ACE1-Ang II-AT1R axis. This could also be a significant contributing factor to the sustained vasoconstriction in the pulmonary vasculature of CKD-PH model rats.

In order to investigate the potential mechanism of RAAS in the development of CKD-PH, metabolomics, a innovative, comprehensive diagnostic tool in clinical and biomedical research was applied. This emerging method has been widely used for analysis of metabolite patterns in CKD rat model. A study indicated that the axis of erythrocyte-specific sphingosine kinase 1 (eSphK1)/Sphingosine 1-phosphate (S1P) has been found to inhibit AngⅡ-induced CKD^52^. Another study reported that 291 serum differential metabolites were identified in a CKD rat model, with polyamine and glycine-conjugated metabolites being associated with the cause and severity of CKD^53^. Enrichment of trimethylamine-N-oxide (TMAO) was also discovered in the serum of 223 CKD patients^54^. Furthermore, linaclotide, an inhibitor of TMAO, was shown to lower uremic toxins (BUN, CCr) and fibrosis of the kidney in a 5/6 nephrectomy-induced CKD mouse model^55^. By applying the same metabolomics profiling methods, ketoacid-stimulated reduction of indoxyl sulfate, betaine, choline, and cholesterol was confirmed in an adenine-induced CKD rat mode^56^, and organic anion transporter 1 (OAT1) was found to modulate interorgan communication in a nephrectomy-induced CKD rat model^57^.

In this manuscript, by integrating metabolomics data, we identified abnormal expression of metabolites in 5/6 Nx group, the metabolites with relative higher expression level are listed as follows: diacylglycerol (DAG), sphinganine, xanthurenic acid, o-propanoylcarnitine, β-D-glucuronoside, N-acetyl-L-phenylalanine, p-hydroxy phenylacetylglycine, indoxyl, 5-hydroxy indoleacetyl-glycine, n-Succinyl-L-glutamate 5-semialdehyde, salicyluric acid, cinnamic acid, 5-methyl-2-furaldehyde, indolin-2-one, alpha-methylstyrene, 3-carbamoyl-2-phenylpro-pionic acid, and dopamine quinone. In particular, DAG was found to have a progressive increase in group continuously given 5/6 Nx and high-salt diet.

Recent report indicated that DAG was implicated in AngⅡ-modified production of aldosterone^58^. As a lipid second messenger, DAG exerted an influence on the production of aldosterone by regulating protein kinase C and calcium homeostasis^59^. Arachidonate and hydroxyeicosatetraenoic acids (HETEs), which were metabolizable by DAG in glomerulosa cells, enhanced the syntheses of aldosterone via regulating Ca^2+^ homeostasis ^60^. What’s more, inhibition of DAG blocked AngⅡ-induced generation of aldosterone^61^. Incidentally, aldosterone was demonstrated contributing to the process of pulmonary hypertension. Aldosterone was increased in MCT-induced rat PH model, and then stimulated the expression of aquaporin 1(AQP1) and proliferation of PASMCs. The antagonists of aldosterone, spironolactone (SP), had a definite therapeutic effect on MCT-induced PH rat model^62^. Combination treatment of SP and ambrisentan was superior to ambrisentan monotherapy in reducing poor exercise tolerance, B-type natriuretic peptide (BNP) and WHO functional class in patients with pulmonary arterial hypertension^63^. The increase of aldosterone was related to the deteriorative hemodynamic measurements and pulmonary vascular remodeling in MCT and SU-5416/hypoxia-induced pulmonary hypertension, and SP reversed this effect^64^. SP exert the anti-inflammatory effect in MCT-induced PH rat model by down-regulating the expression of xeroderma pigmentosum group B complementing protein (XPB) ^65^.

DAG is one of the metabolisms of fatty acid, which play a role in pulmonary hypertension and right heart failure^66^. it’s reported that DAG was a key factor in modulating vascular tone of pulmonary artery by a mechanism involving with Transient receptor potential canonical channel 6 (TRPC 6) and cellular calcium homeostasis^67–70^. AngⅡ-induced constriction of human and rat pulmonary arteries was blocked by suppressor of DAG^71^. Besides, DAG enhanced mechanosensitive cation currents and cellular Ca2+ influx of human pulmonary arterial endothelial cells (PAECs) mediate TRPC6^72^. According to the KEEG database, we found DAG was associated with synthesis and secretion of aldosterone, indicating RASS might exert an influence on CKD-PH via DAG.

In conclusion, by using the combination treatment of 5/6 nephrectomy and high-salt diet in rats, we successfully established CKD associated PH rat model. This animal model was characterized by the impairment of kidney, overload of pulmonary artery pressure and deteriorated heart function. Notably, during the time course of 5/6 nephrectomy and high-salt diet treatment, significant increased pressure of PA was observed in 14 weeks with the absent of pulmonary vascular remodeling. We confirmed the activation of RAASS in this CKD-PH rat model. By untargeted metabolomics, we then discovered DAG was raised with time.

In summary, we strongly believe that AngⅡand DAG increased as the molding time prolongs, causing the accumulation of aldosterone and constriction of PA, ultimately leading to CKD associated PH. Our findings revealed a novel insight of CKD associated PH and might be useful for the treatment of this subtype of PH.

### Perspectives

This study revealed that the formation of pulmonary arterial hypertension in CKD-PH rats is characterized by an increase in pulmonary artery pressure. There is an elevation in serum vasoconstrictive substances, a downregulation of pulmonary endothelial ACE2 expression, and a potential mechanism for the elevation of pulmonary artery pressure may be the increase in vasoconstrictive substances found in serum metabolites. Inhibiting the disruption of the RAAS system and metabolic abnormalities caused by chronic kidney disease may be an effective strategy for intervening in CKD-PH.

## Acknowledgements

None.

## Contributions

Q. Jiang performed functional and molecular biological experiments, analyzed data, prepared figures, and drafted article; Q. Yang, C. Zhang, C. Hou, W. Hong, M. Du, X Shan, X. Li and D. Zhou performed animal experiments. D. Wen, Y. Xiong, K. Yang, Z. Lin, J. Song, Z. Mo, H. Feng, Y. Xing and X. Fu performed molecular biological experiments. C. Liu, F. Peng, B. Li and W. Lu contributed to technical support and reagents. J. Wang initiated and provided critical suggestions to the project and revised the article; Y. Chen initiated the project, designed the experiments, wrote, and revised the article; J. X.-J. Yuan provided critical suggestions in the design of the project; All authors approved for the submission of the article.

## Non-standard Abbreviations and Acronyms

ACE: Angiotensin converting enzyme
ALD: Aldosterone
AQP1: Aquaporin 1
BNP: B-type natriuretic peptide
CKD: Chronic kidney disease
CO: Cardiac output
COPD: Chronic obstructive pulmonary disease
CTEPH: Chronic thromboembolic pulmonary hypertension
DAG: Diacylglycerol
DBP: Diastolic blood pressure
EF: Ejection fractions
eSphK1: Erythrocyte-specific sphingosine kinase 1
FC: Fold change
HETEs: Hydroxyeicosatetraenoic acids
IPAH: Idiopathic pulmonary arterial hypertension
KEGG: Kyoto Encyclopedia of Genes and Genomes
mABP: mean arterial blood pressure
OAT1: Organic anion transporter 1
PADN: Percutaneous pulmonary artery denervation
PAECs: Pulmonary arterial endothelial cells
PASMCs: Pulmonary arterial smooth muscle cells
PAT/PET: Pulmonary acceleration time/pulmonary ejection time
PH: Pulmonary hypertension
RAAS: Renin-angiotensin-aldosterone system
ROC: Receiver-operating-characteristic
RVH: Right ventricular hypertrophy
RV/(LV+S): right ventricular/(left ventricular + septum)
RVSP: Right ventricular systolic pressure
SBP: Systolic blood pressure
SNS: Sympathetic nervous system
SP: Spironolactone
SV: Stroke volume
S1P: Sphingosine 1-phosphate
TMAO: Trimethylamine-N-oxide
TRPC6: Transient receptor potential canonical channel
XPB: Xeroderma pigmentosum group B complementing protein

## Novelty and Relevance

### What Is New?

We established a rat model of chronic kidney disease with pulmonary arterial hypertension (CKD-PH) by performing 5/6 nephrectomy surgery and administering a high-salt diet. The pathological features of this model primarily manifest as an elevation in pulmonary artery pressure.

### What Is Relevant?

Inhibiting the disruption of the RAAS system and metabolic abnormalities caused by chronic kidney disease may be an effective strategy for intervening in CKD-PH.

